# Long-range gene regulation network of the *MGMT* enhancer in modulating glioma cell sensitivity to temozolomide

**DOI:** 10.1101/2020.11.03.367268

**Authors:** Anshun He, Bohan Chen, Jinfang Bi, Wenbin Wang, Jun Chen, Yuyang Qian, Tengfei Shi, Zhongfang Zhao, Jiandang Shi, Hongzhen Yang, Lei Zhang, Wange Lu

**Author notes:** These authors contributed equally to this work (Anshun He, Bohan Chen). Corresponding authors: Wange Lu and Lei Zhang.

## Abstract

Acquired resistance to temozolomide (TMZ) is a major obstacle in glioblastoma treatment. MGMT, a DNA repair protein, and the methylation at its gene promoter, plays an important role in TMZ resistance. However, some evidences have suggested a MGMT-independent mechanisms underlying TMZ resistance. Here, we used *MGMT* enhancer as a model and discovered that its deletion in glioma cells of low *MGMT* expression induced increased sensitivity to temozolomide. Analysis of a combination of RNA-seq and Capture Hi-C further suggested multiple long-range target genes regulated by the *MGMT* enhancer and that interactions may play important roles in glioma cell sensitivity to TMZ. This study reveals a novel mechanism of regulation of TMZ sensitivity in glioma cells.

## Introduction

Approximately 80% of primary tumors in the central nervous system are malignant glioma (1), and >50% of those are diagnosed as glioblastoma (GBM), the most aggressive glioma. Despite treatment advances, glioblastoma prognosis is unsatisfying, and median survival time of patients is only 14~17 months (2).

Temozolomide (TMZ) is a reagent frequently used in the clinic as chemotherapy for GBM (3,4), due to its high bioavailability and relatively low side effects. However, development of chemo-resistance to TMZ significantly decreases its efficacy and remains a major treatment obstacle. Thus it is proposed that increasing glioma cells’ sensitivity to TMZ could improve prognosis of patients with GBM. Several previous studies report that O-6-methylguanine-DNA methyltransferase (MGMT) promotes TMZ resistance in glioma (5,6) by removing cytotoxic O-6-methylguanine-DNA lesions generated by TMZ. Other factors associated with TMZ resistance include poly (ADP-ribose) polymerase (7), ALDH1A1 (8), and P4HB (9). Moreover, Gaspar and colleagues report that phosphoinositide 3-kinase-mediated *HOXA9/HOXA10* expression is associated with MGMT-independent TMZ-resistance in glioblastoma cells (10). In addition, patients whose tumor cells show no apparent expression of *MGMT* are susceptible to TMZ resistance (11), indicating that other factors may also underlie TMZ resistance. These findings are supported in part by the fact that MGMT inhibitors show little effect against glioma in clinical trials (12,13).

A previously identified *MGMT* enhancer was reported to be associated with methylation of *MGMT* promoter and expression of *MGMT* (14,15). However, inconsistency of methylation level of *MGMT* promoter and *MGMT* expression were also found in GBM (16–18). Thus, *MGMT* expression may be regulated by other enhancers or factors (19), or the *MGMT* enhancer may regulate genes other than *MGMT*, which may contribute to its role in TMZ resistance.

High-order chromatin structure studies have revealed that regulatory elements like enhancers can regulate long-range gene expression (20–23). Thus we asked whether the *MGMT* enhancer may not only regulate MGMT but also other genes through long-range interactions and in so doing alter TMZ sensitivity of glioma cells. Here, we report that in glioma cells of low MGMT expression a *MGMT* enhancer regulates glioma cell sensitivity to TMZ through long-range regulation of multiple target genes, including *MKI67*, providing a novel regulatory mechanism underlying glioma cells TMZ sensitivity.

## Results

### *MGMT* enhancer deletion in glioma cells showing low MGMT expression increases their sensitivity to TMZ

To determine whether the *MGMT* enhancer is associated with glioma cell sensitivity to TMZ we used the CRISPR/Cas9 system to delete the *MGMT* enhancer region in U251 glioma cells, whose *MGMT* expression is low (24). Two enhancer KO lines (KO-1, lacking 874bp, and KO-2, lacking 573bp of the enhancer region) were generated (Fig. 1A, Supplementary Fig. S1). Wickstrom et al. previously reported Western blot data showing no MGMT expression in five of six typical glioma lines, including U251, and proposed this deficit was due to *MGMT* promoter methylation (24). Our RT-qPCR analysis confirmed low *MGMT* transcription levels in wild-type (WT) U251 cells treated 72 hours with TMZ (1mM) or control vehicle (Supplementary Fig. S2A). Moreover, *MGMT* enhancer deletion did not significantly alter *MGMT* transcription in either KO line (Supplementary Fig. S2B). We then treated both enhancer KO lines and WT U251 cells with TMZ (1mM) for 72 hours and assessed potential cytotoxicity by performing lactate dehydrogenase (LDH) and caspase 3/7 activity assays (Fig. 1B). Cytosolic LDH is released to the medium by dying cells, and caspase 3/7 activity is an apoptotic marker. Compared with TMZ-treated WT U251 cells, LDH levels in culture medium were significantly increased by 2-3 fold (after 24h) and 4-5 fold (after 48h) in *MGMT* enhancer KO glioma cells after TMZ treatment (Fig. 1C,D). Caspase 3/7 activities in KO cells also significantly increased (~3-fold) relative to WT U251 cells after TMZ treatment at 48h and 72h (Fig. 1E,F). Given that MGMT is not detectable in U251 cells treated with or without TMZ (1mM, 72h), these data suggest that the *MGMT* enhancer modulates glioma cell TMZ sensitivity through genes other than *MGMT*.

**Figure 1.**
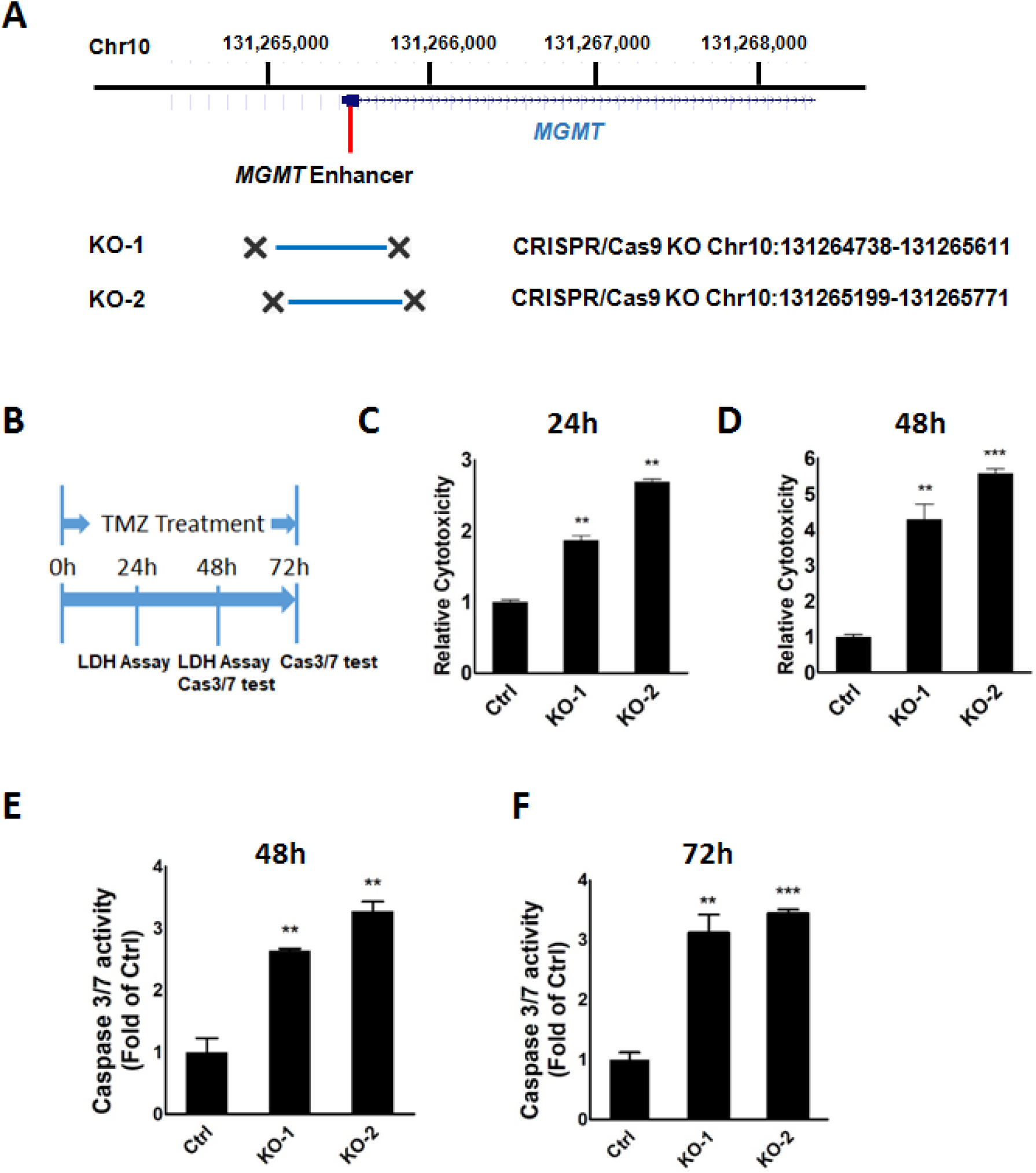
*MGMT* enhancer deletion increases glioma cell sensitivity to TMZ. A, Deletion of the *MGMT* enhancer in KO-1 (deleted region: hg19_ Chr10: 131264738-131265611) and KO-2 (deleted region: hg19_Chr10: 131265199-131265771) lines. B, Schematic showing time frame for LDH and Caspase 3/7 assays in U251 glioma and *MGMT* enhancer KO glioma lines treated with TMZ (1mM) as indicated. C, D, LDH release assays were used to detect cytotoxicity in wild-type U251 (Ctrl) and *MGMT* enhancer knock-out (KO-1, KO-2) U251 lines after 24h or 48h of TMZ (1mM) treatment. Data represents means ± S.E.M. of three independent experiments; **p < 0.01, ***p < 0.001 compared with control. E,F, Caspase 3/7 activities were assayed in wild-type (Ctrl) or *MGMT* enhancer knock-out (KO-1, KO-2) U251 lines treated 48h or 72h with TMZ (1mM). Data represents means ± S.E.M. of three independent experiments; **p < 0.01, ***p < 0.001 compared with control.

### Long-range interactions of the *MGMT* enhancer with target genes

Studies of high-order chromatin structure indicate that enhancers can regulate long-range targets though 3D genome structure. Hi-C data analysis in glioma cells treated with or without TMZ showed that TMZ treatment altered high-order chromatin structure (Fig. 2A). Specifically, a Z-score map showed global changes in interaction frequencies in the *MGMT* enhancer region (Fig. 2A), suggesting that long-range interactions are associated with drug sensitivity. To investigate potential *MGMT* enhancer target genes, we performed Capture Hi-C in U251 glioma cells. We used a 1.8 kb region containing the *MGMT* enhancer as a “bait” region (Fig. 2B) and performed Capture Hi-C based on a previously reported protocol (25, 26). Interactions between the bait region and target chromatin regions from two biological replicates are depicted as curves in Circos plots shown in Fig. 2C,D (27). *In cis* interacting sites comprised 73.3% and 69.8% of respective biological replicates. When we mapped sites to the human genome, we identified 89 target genes, 84 *in cis* and 5 are *in trans*, which physically contacted with the *MGMT* enhancer in both biological replicates. RNA-seq data in WT and KO-1 and KO-2 cells revealed global transcriptional changes in enhancer KO relative to WT U251 cells (Fig. 3A). Gene Ontology (GO) analysis of significantly changed genes (q <0.05) between enhancer KO and WT U251 cells identified focal adhesion as the top altered signaling pathway (Fig. 3B), an observation consistent with a previous report that focal adhesion activity is associated with glioma cell TMZ sensitivity (28). We then performed combination analysis of RNA-seq and Capture Hi-C to confirm direct target genes potentially regulated by the *MGMT* enhancer. Genes with a positive signal in Capture Hi-C data are shown in Fig. 3C. Moreover, Fig. 3C shows the names of genes (*KIAA1549L, SH3XPD2A, ADAM12* and *MKI67*) with significant altered mRNA expression in the *MGMT* enhancer KO relative to WT U251 cells (WT/KO fold change > 1.5; q value < 0.05). *SH3XPD2A, ADAM12* and *MKI67* were *in cis* targets and *KIAA1549L in trans* (on chromosome 11). RT-qPCR analysis showed decreased *KIAA1549L, SH3XPD2A, ADAM12* and *MKI67* transcript levels in enhancer KO lines relative to wild-type U251 cells, with or without TMZ treatment (Fig. 3D). Capture Hi-C counts at *MKI67* and *ADAM12* loci were significantly higher than *KIAA1549L* or *SH3XPD2A* (Fig. 3C), suggesting strong interactions at the *MKI67* and *ADAM12* loci.

**Figure 2.**
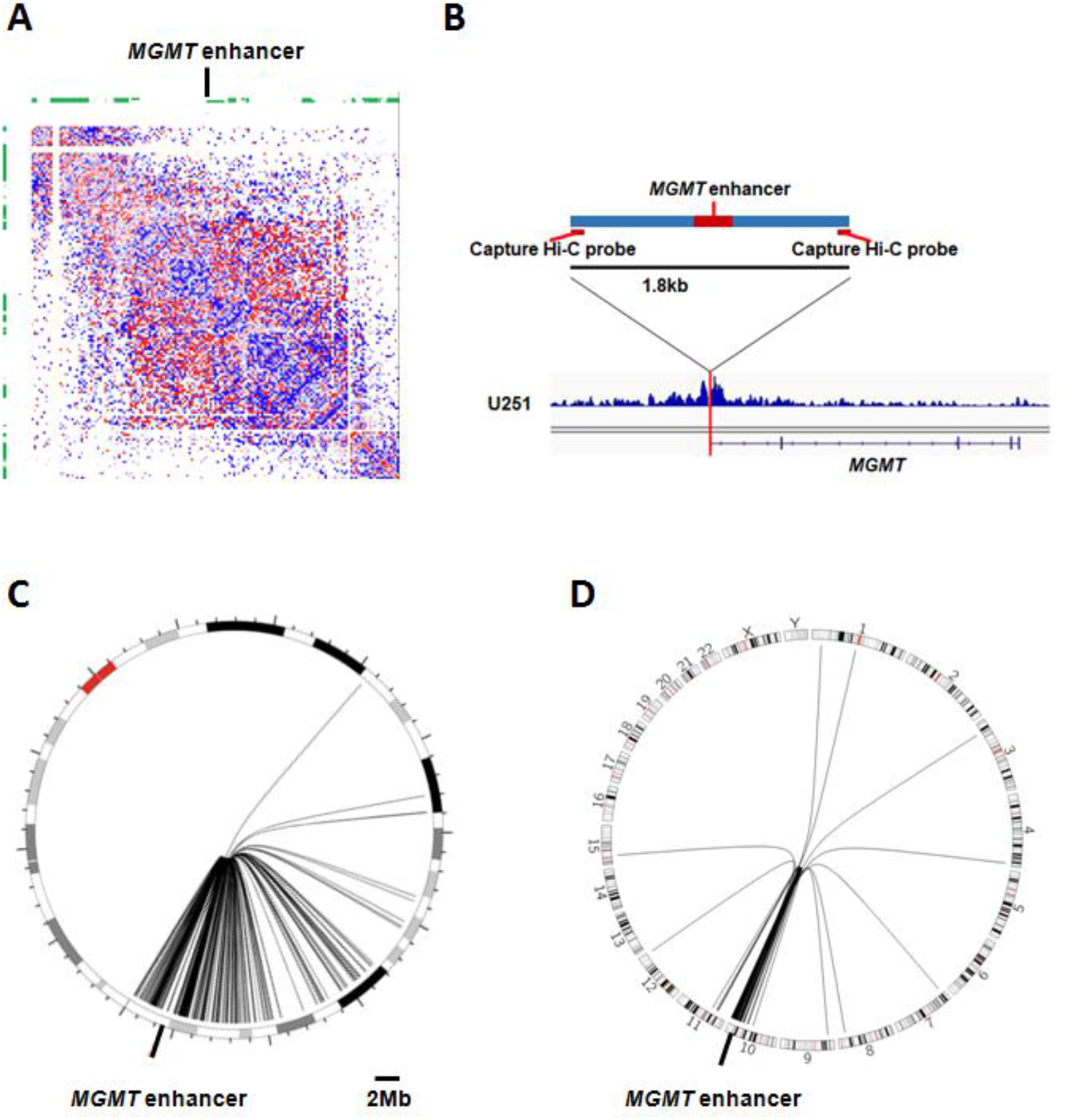
*MGMT* enhancer: intra- and inter-chromosomal interactions. A, Z-score difference maps for a chromatin region ± 3Mb from the *MGMT* regulatory element binned at 40kb resolution. Increased (red) and decreased (blue) frequency of chromatin interactions seen in TMZ-treated (1mM, 72h) U251 glioma cells compared with untreated control cells are shown. B, 1.8kp Capture Hi-C bait and Capture Hi-C signal peaks near the bait region. C, Circos plot showing same intra-Chr10 interactions from two biological replicates, indicated by curves extending from the bait region. D, Circos plot showing same genome-wide intra- and inter-interactions from two biological replicates, indicated by curves extending from the bait region.

**Figure 3.**
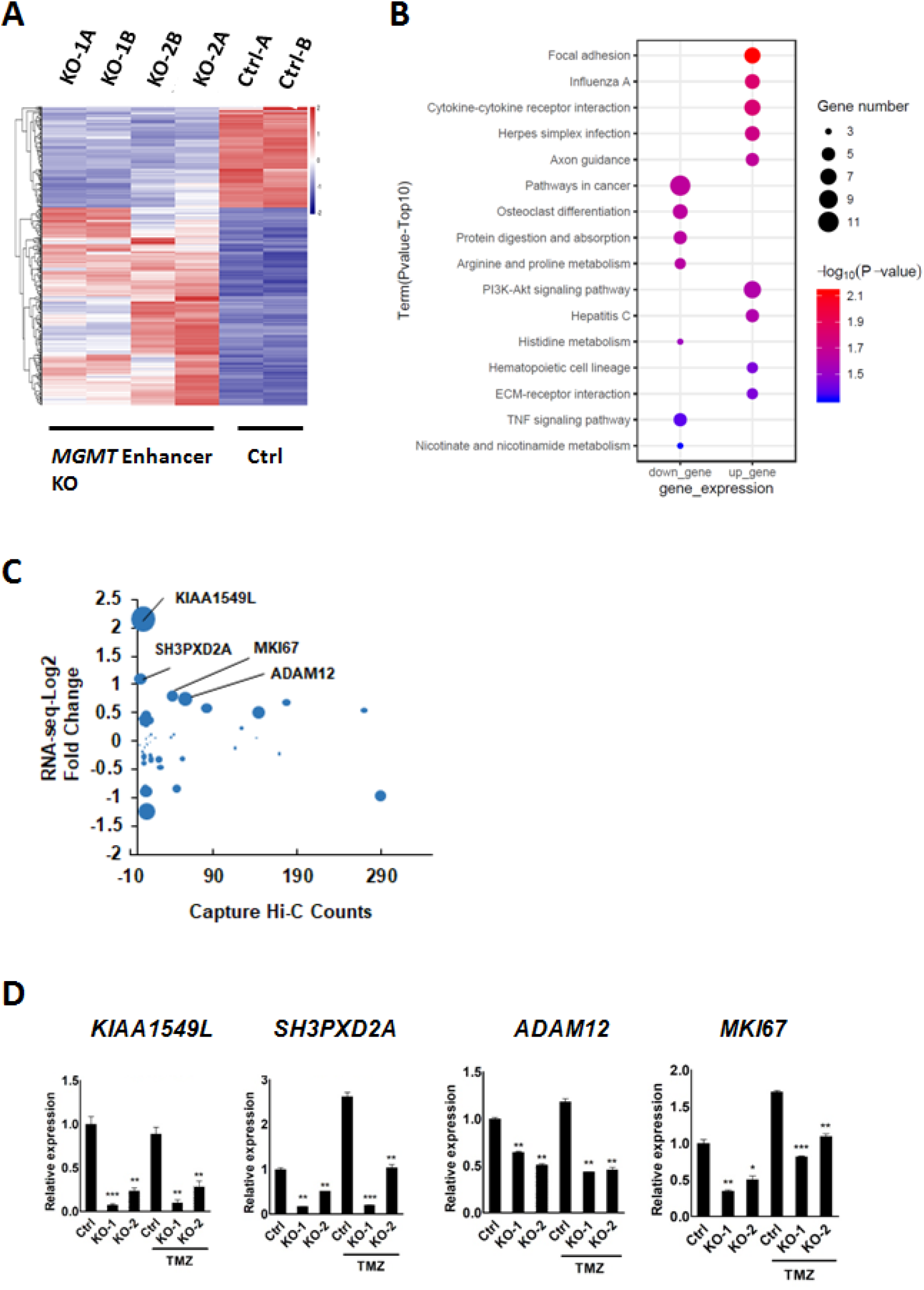
Analysis of RNA-seq and Capture Hi-C data. A, Changes in gene expression in *MGMT* enhancer knock-out (KO) and wild-type (Ctrl) glioma lines. Heatmap shows clustering of differentially expressed genes in enhancer KO and Ctrl glioma cells. KO-1A and B, KO-2A and B, Ctrl-A and B refer to two biological replicates. B, GO-KEGG pathway analyses of differentially expressed genes (p<0.05). C, Combination analysis showing fold-change in gene expression based on RNA-seq data and Capture Hi-C signals. Potential target genes reproducible in two biological replicates of Capture Hi-C are shown on the plot. Four genes showing significant expression changes (adjusted p value < 0.05, and fold-change > 1.5 or < −1.5) are indicated by name. D, RT-qPCR analysis of mRNA (independent mRNA library) levels of indicated potential target genes in Ctrl and *MGMT* enhancer KO cells treated with or without TMZ (1mM). Data represents means ± S.E.M. of three independent experiments. *p<0.05, **p<0.01, ***p<0.001 compared with corresponding Ctrl cells.

### *MKI67* upregulation decreases TMZ sensitivity in glioma cells lacking the *MGMT* enhancer

To determine whether *ADAM12* and *MKI67* are associated with increased sensitivity of glioma cells TMZ, we overexpressed both genes separately in KO-1 and KO-2 lines. Following transfection of glioma cells with *ADAM12* expression constructs, we verified the overexpression by RT-qPCR (Fig. 4A). However, we observed no significant change in TMZ sensitivity relative to control cells in either KO-1 or KO-2 lines following ADAM12 overexpression based on either LDH (Fig. 4B) or caspase 3/7 (Fig. 4C) assays. We then overexpressed *MKI67* in WT U251 and enhancer KO cells using the CRISPRa system and observed 3-4-fold increases in *MKI67* transcript levels in all engineered lines (Fig. 5A). Moreover, we observed a significant decrease in TMZ sensitivity, as indicated by decreased LDH release, in *MK167*-overexpressing *MGMT* enhancer KO cells relative to control KO-1 and KO-2 cells following 48 hours of TMZ (1mM) treatment. Accordingly, *MKI67* overexpression induced a significant decrease in caspase 3/7 activities in *MGMT* enhancer KO cells treated 72h with TMZ (1mM) (Fig. 5C). We conclude that *MKI67* upregulation rescues increased TMZ sensitivity of glioma cells lacking the *MGMT* enhancer, and that regulation of *MKI67* by *MGMT* enhancer may play an important role in modulating glioma cells’ TMZ sensitivity. The long-range interactions of *MKI67* and *MGMT* enhancer was confirmed by a 3C assay (Fig. 5D,E).

**Figure 4.**
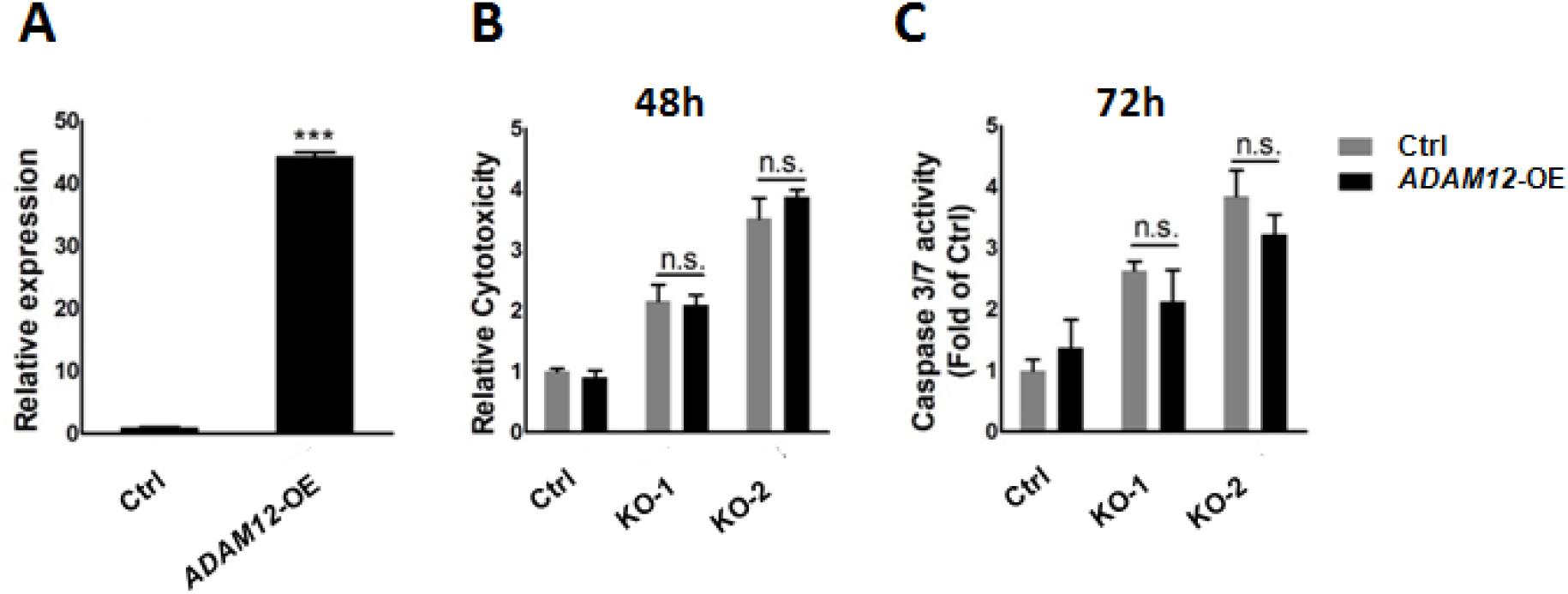
*ADAM12* overexpression does not significantly rescue increased TMZ sensitivity in *MGMT* enhancer KO glioma cells. A, RT-qPCR verification of *ADAM12* overexpression (*ADAM12*-OE) efficiency. Data represents means ± S.E.M. of three independent experiments. ***p<0.001 compared with Ctrl. B, LDH release was tested in control U251 glioma cells and *MGMT* enhancer KO glioma cells with overexpressed *ADAM12* (*ADAM12-OE*). LDH release (shown as “Relative Cytotoxicity”) was assayed after 48h of TMZ (1mM) treatment. Data represents means ± S.E.M. of three independent experiments. C, Caspase 3/7 activities were assayed in control U251 glioma and *MGMT* enhancer KO glioma cells overexpressing *ADAM12* (*ADAM12-OE*). Caspase 3/7 activities were assayed after 72 h of TMZ (1mM) treatment. Data represents means ± S.E.M. of three independent experiments.

**Figure 5.**
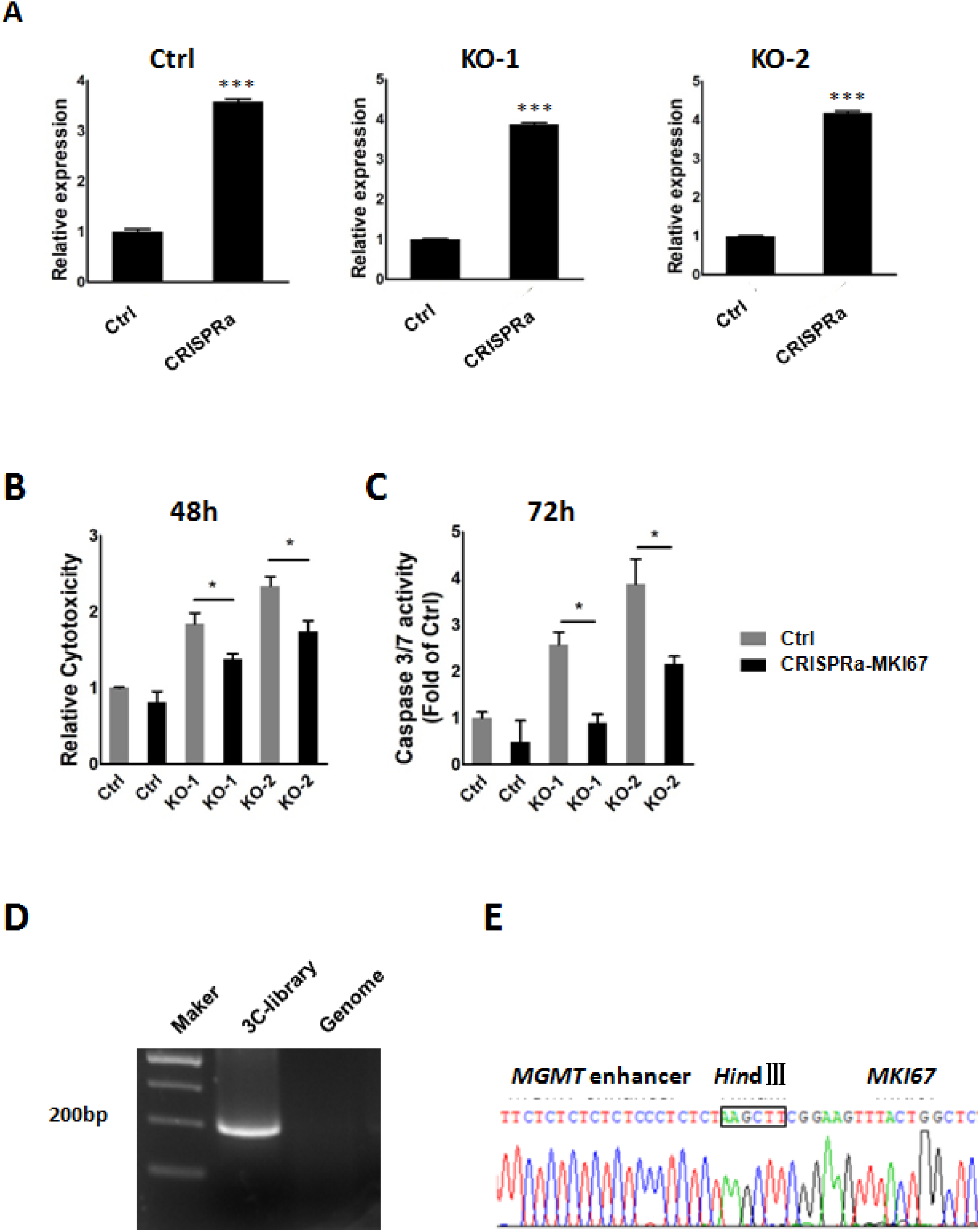
*MKI67* overexpression rescues increased TMZ sensitivity in *MGMT* enhancer KO glioma cells. A, RT-qPCR verification of MKI67 overexpression (MKI67-OE). Data represents means ± S.E.M. of three independent experiments. ***p<0.001 compared with Ctrl. B, LDH release was assayed in control U251 glioma and MGMT enhancer KO glioma cells overexpressing MKI67 (MKI67-OE). LDH release was evaluated after 48h of TMZ (1mM) treatment. Data represents means ± S.E.M. of three independent experiments. *p<0.05 compared with corresponding Ctrl. C, Caspase 3/7 activities were assayed in control U251 glioma and MGMT enhancer KO glioma cells overexpressing MKI67 (MKI67-OE). Caspase 3/7 activities were evaluated after 72h of TMZ (1mM) treatment. Data represents means ± S.E.M. of three independent experiments. *p<0.05 compared with corresponding Ctrl. D,E MGMT enhancer/MKI67 interactions were verified by 3C-PCR and sequencing. 3C-library: 3C PCR products from the DNA fragments in the 3C library. Genome: Genomic DNA as the PCR template. Forward and reverse PCR primers were designed based on MGMT enhancer and MKI67 gene regions.

## Discussion

Development of chromosome conformation capture technologies has enabled study of long-range chromatin interactions (25, 29–32). Moreover, several previous studies report long-range regulation of target genes by gene regulatory elements, including enhancers (22, 23, 33) and repressive regulatory elements (20, 34). Here we report that in glioma cells, a *MGMT* enhancer can regulate expression of the multiple target genes including *MKI67*, which is located ~1.33Mb away the enhancer locus through long-range chromatin interactions and modulate glioma cell TMZ sensitivity. This study reveals novel mechanism underlying glioma cell drug sensitivity. Previous researches have reported that enhancers or repressive elements can regulate long-range target genes in cancer cells. For example, a recent study showed that in prostate cancer an active enhancer regulates several target genes through long-range intra- and inter-chromatin interactions to alter tumor cell proliferation and invasion (23). Another study showed that a prostate cancer-specific enhancer may regulate *SOX9* through a 1Mb chromatin loop (35). However, few studies investigate the long-range regulation network of *MGMT* enhancer besides its function on regulating *MGMT*. Our research identifies long-range gene regulation network of the *MGMT* enhancer and indicates that long-range gene regulation by the *MGMT* enhancer can modulate glioma cell sensitivity to TMZ, which is MGMT-independent mechanism underlying TMZ resistance.

Combination analysis using RNA-seq data and Capture Hi-C data identified 89 target genes that might be regulated by the *MGMT* enhancer. *MKI67* and *ADAM12* showed significant strong interactions with MGMT enhancer compared with other target genes, and *MKI67* overexpression induced a significant decrease in TMZ sensitivity,suggesting *MKI67* may play an important role in the gene network regulated by the *MGMT* enehancer. The *MKI67* gene encodes Ki67, a proliferation marker often monitored in cancer cells, including glioma (36–38). High Ki67 levels are associated with increased proliferation and, in the case of brain cancer, poor prognosis (39). Analysis of survival of glioma patients also indicates that low *MKI67* expression is associated with longer patient survival time (Supplementary Fig. S3). Ki67 reportedly prevents chromosomes from collapsing into a single chromatin mass and enables independent chromosome motility (40). Booth and colleagues reported that Ki67 may function in nucleolar segregation between daughter cells (41). In addition, Ki67 is reportedly associated with ribosomal RNA (rRNA) synthesis (42). Sobecki et al found that Ki-67 likely modulates transcription in cancer cells by regulating heterochromatin organization (43). These studies overall suggest that manipulating Ki67 activity may alter cell cycling and modulate effects of TMZ treatment in glioma cells (44). Data presented here strongly suggests that *MKI67* is regulated by the *MGMT* enhancer and that *MKI67* downregulation following enhancer deletion increases glioma cell sensitivity to TMZ.

Data in this study also indicate that *ADAM12* might be a candidate target of the *MGMT* enhancer, and others have reported that ADAM12 promotes cell proliferation and migration (45, 46). However, we observed no significant change in glioma cell TMZ sensitivity in *MGMT* enhancer KO cells following *ADAM12* overexpression. Overall, the regulation network likely governed by the *MGMT* enhancer may include several target genes and pathways whose functions should be further addressed in future studies.

In summary, our study shows that a *MGMT* enhancer regulates glioma cell sensitivity to TMZ by long-range regulation of multiple genes. Modifying chromatin structure to change long-range regulations using the CRISPR system or artificial zinc fingers has been reported recently (47–49). Our study suggests that changing interactions between the *MGMT* enhancer and target genes might be a novel way of increasing glioma cell sensitivity to TMZ.

## Experimental procedures

### Cell culture

The GBM cell line U251 was obtained from the National Infrastructure of Cell Line Resource (China). U251 glioma cells were incubated at 37 °C with 5% CO_2_ and cultured in Dulbecco’s Modified Eagle Medium (GIBCO), supplemented with 10% fetal bovine serum (FBS) (BI) and 1% penicillin/streptomycin (GIBCO). All cells were tested for mycoplasma contamination monthly and found to be mycoplasma-free.

### CRISPR/Cas9-mediated deletions

U251 glioma cells were transfected with Cas9 and guide RNA plasmids that target the MGMT enhancer core region (hg19_Chr10:131265544-131265602). Guide RNAs were designed using the MIT CRISPR Design website (http://crispr.mit.edu). Only high-score guide RNAs (score>85) were used to minimize potential off-target effects. Guide RNA sequences are listed in Table S1. *MGMT* enhancer KO clones were genotyped by PCR (primers are listed in Table S1), and 2 clones (KO-1 and KO-2) with homozygous deletion of *MGMT* enhancer were used for experiments.

### Lactate dehydrogenase assay

Lactate dehydrogenase (LDH) in culture medium was evaluated using a LDH cytotoxicity kit (Promega) following the manufacturer’s instructions. Briefly, cells were cultured in 96-well plates and after cells were treated with TMZ or transfected, the culture medium was collected and LDH activity was assayed. Levels of released LDH in experimental groups were measured as OD values by using a microplate reader and calculated as a percentage of the total amount (the positive control is described in the manufacturer’s instructions).

### Caspase 3/7 activity measurement

A caspase - Glo 3/7 kit (Promega) was used to evaluate caspase 3/7 activity based on the kit protocol. Cells were harvested and incubated with 100μL caspase - Glo 3/7 reagent for 1 hour on a rotary shaker in the dark. Sample luminescence was measured at 485/530 nm using a microplate reader. Relative caspase 3/7 activity was calculated to evaluate fold-changes in samples either drug-treated or transfected relative to controls.

### *ADAM12* overexpression

U251 glioma were transiently transfected with *ADAM12* overexpression plasmids (Sino Biological) using Lipofectamine 3000 (Thermo Fisher Scientific) according to the manufacturer’s instructions.

### *MKI67* overexpression

*MKI67* was overexpressed in glioma cells using the CRISPR-activation (CRISPRa) system as reported (50). In brief, a stable U251 CRISPRa cell line was generated using lentiMPH v2 plasmid (Addgene). Lentiviral particles were generated in 293T cells using pMDG.2 (Addgene) and psPAX2 (Addgene) packaging plasmids. U251 cells were transduced for 24h and selected in 200 mg/ml Hygromycin (Invitrogen) for 5 days. CRISPRa sgRNAs targeting the *MKI67* promoter were designed using GPP Web Portal (Broad institute; https://portals.broadinstitute.org/gpp/public/analysis-tools/sgrna-design) and ordered from Sangon Biotech (Table S2). Sequences were subsequently cloned into lentiSAM v2 plasmid (Addgene) using *Bsm*BI (New England Biolabs), packaged and cells transduced as described above. Transduced U251 CRISPRa cells were selected in 3 mg/ml blasticidin (Solarbio).

### RNA-seq

Total RNA was extracted from U251 glioma cells or enhancer KO cells. Barcoded RNA-seq libraries were sequenced as 150bp paired-end reads using the Illumina HiSeq 4000 platform. Reads were mapped to the human reference genome (hg19) using HISAT (51) with a GENCODE GTF file supplied as gene model annotations. HTSeq (52) was used to quantitate transcript abundance for each gene. DESeq2 (53) was used to perform normalization and regularized log transformations on read counts.

### Survival curves

Survival data relevant to glioma patients were obtained from the TCGA database. Patient samples were divided into two groups based on expression level of *MKI67* as indicated in Results. Glioma samples with gene expression levels higher or lower than the median were classified respectively as “high expression” or “low expression” groups. A P value < 0.0001 indicates a significant survival difference between groups.

### Capture Hi-C and Hi-C

Capture Hi-C and Hi-C were performed following previously reported protocols (25,26). For each sample, two biological replicates were assessed with 1×10 U251 cells. The Hi-C library was generated with *Dpn*II digestion and then sheared to 200-300bp fragments by sonication. Then the library or pre-capture library was prepared using the NEBNext DNA library kit (New England BioLabs) according to the manufacturer’s instructions. Biotinylated probes and streptavidin beads (Thermo Fisher Scientific) were used to enrich “bait” regions and linked chromatin fragments by two rounds of hybridization-capture approach. Capture Hi-C libraries were sequenced as 150bp paired-end reads using the Illumina HiSeq X Ten platform. Capture Hi-C and Hi-C data were analyzed based on a pipeline proposed in previous studies (25,26,54).

Two biotinylated DNA oligonucleotides were designed for both ends of the region containing the *MGMT* enhancer, with the following sequence (5’ to 3’):

Biotin-GATCCTGCTCCCTCTGAAGGCTCCAGGGAAGAGTGTCCTCTGCTCCCTCCGAAGGCTCCAGGGAAGGGTCTGTCCTCTTAGGCTTCTGG
Biotin-TGCTCTCAGTTGCTTCAGCTGAGTAGCTGGCTTTCTGTCCTGGAAAGCAGACTTTGTACATGTGTGTGCAACCTATGCCTGCTGAGATC

### Gene Ontology (GO) analyses

GO and KEGG pathway analyses were performed using the DAVID knowledgebase (http://david.abcc.ncifcrf.gov/) (55,56).

### Quantitative real-time PCR

Total RNA was isolated from glioma cells using TRIzol Reagent (Invitrogen) following the manufacturer’s protocol. cDNAs were reverse-transcribed from 5μg total RNA using a PrimerScript RT reagent kit (Takara). RT-qPCR was performed using an SYBR Premix Ex Taq (Takara). *GAPDH* served as an internal reference, and relative expression of target genes was quantified using the 2^-ΔΔCt^ method. Primer sequences used in this study are listed in Table S3.

### 3C assays

1×10 U251 cells were used to prepare 3C libraries according to a previously described protocol (29), and 3C libraries were generated using *Hin*dIII digestion. Interactions between the *MGMT* enhancer region and *MKI67* were detected using nest-PCR. Primer information is listed in Table S4.

### Statistical Analysis

Data represents means ± S.E.M.; statistical analysis was performed using Student’s *t*-test. *p<0.05, **p<0.01, ***p<0.001, ****p<0.0001.

## Data availability

All sequencing data generated in this study have been submitted to the NCBI Gene Expression Omnibus (GEO; https://www.ncbi.nlm.nih.gov/geo/) under accession numbers GSE125629, GSE129476 and GSE125243.

## Acknowledgments

This work was supported by the National Natural Science Foundation of China (Grant No. 31701129, 31530027, 81772687), the Natural Science Foundation of Tianjin City of China (Grant No. 18JCQNJC10100), National Key R&D Program of China (NO.2017YFA0102600) and the Fundamental Research Funds for the Central Universities, Nankai University (63201087). We thank Dr. Lingyi Chen at the College of Life Sciences of Nankai University for providing Cas9 plasmids.

## Authors’ contributions

A.H., L.Z., B.C., J.B., J.C., D.G., YQ., W.W., T.S., Z.Z., J.S., W.A., and F.A. conducted the experiments; L.Z. and W.L. designed the experiments; and L.Z. and W.L. wrote the paper.

## Funding and additional information

National Natural Science Foundation of China (31701129), recipient: L.Z.

National Natural Science Foundation of China (31530027), recipient: J.S.

National Natural Science Foundation of China (81772687), recipient: W.L.

National Key R&D Program of China (2017YFA0102600), recipient: W.L.

Natural Science Foundation of Tianjin City of China (18JCQNJC10100), recipient: L.Z.

Fundamental Research Funds for the Central Universities, Nankai University (63201087), recipient: L.Z.

## Supplementary Figures

**Supplementary Figure S1.**
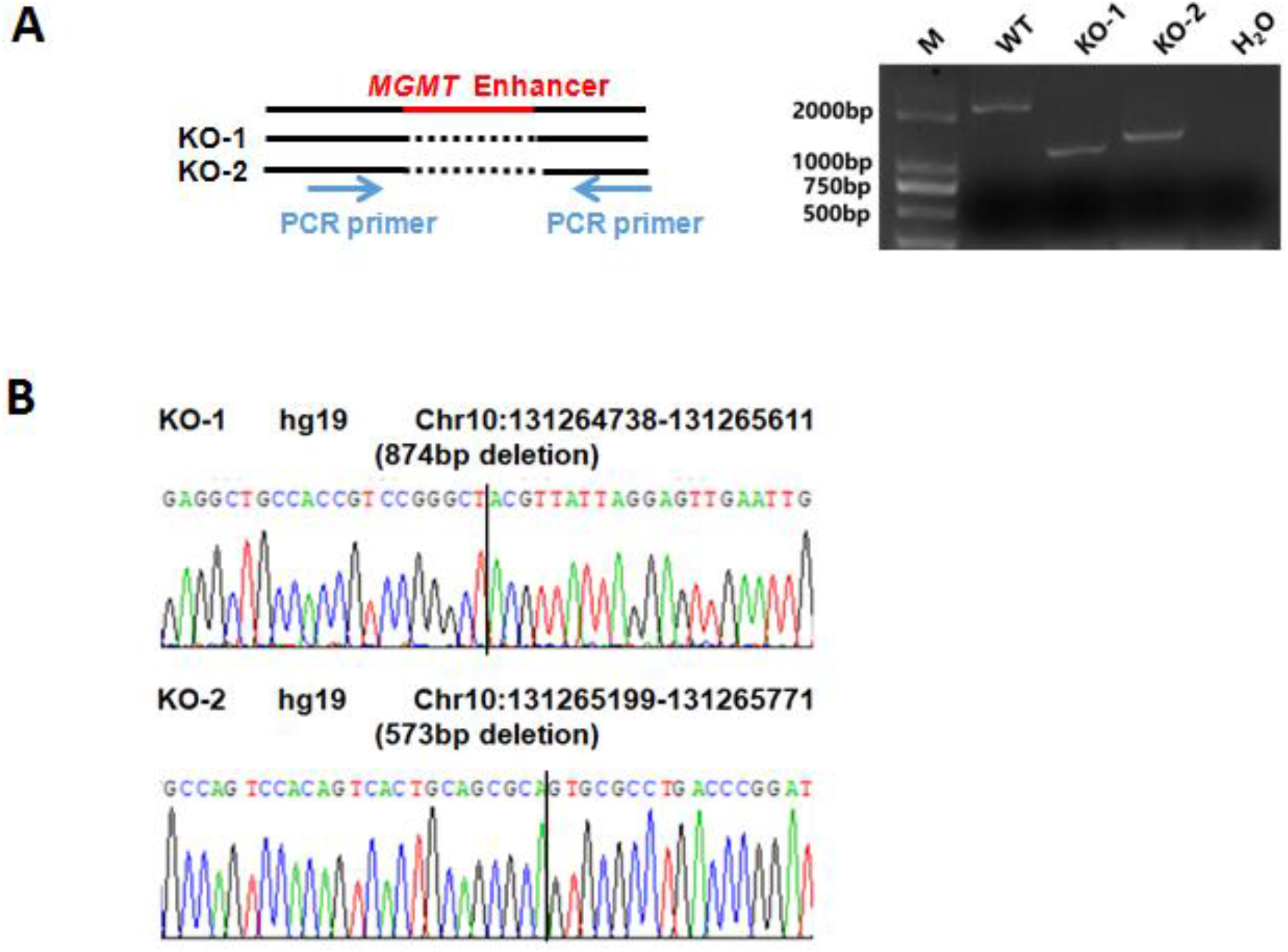
CRISPR/Cas9-mediated deletion of the *MGMT* enhancer. A, PCR analysis used to identify *MGMT* enhancer KO cell lines. The PCR product of the *MGMT* enhancer region is shown in the WT lane. M: marker; H_2_O: negative control. B, Sequencing of the *MGMT* enhancer region in KO cells.

**Supplementary Figure S2.**
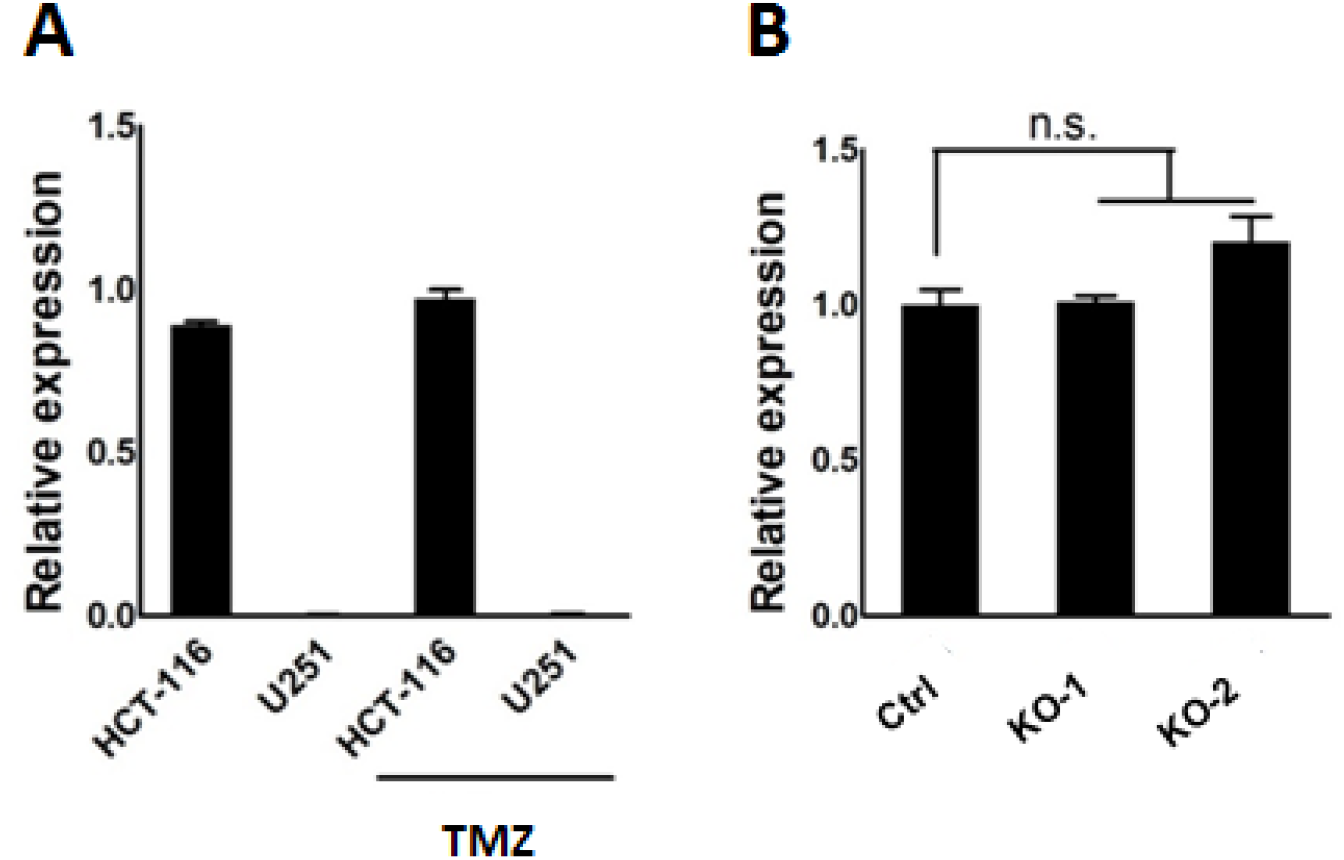
*MGMT* transcript levels in U251 cells and *MGMT* enhancer KO cells. A, RT-qPCR analysis of *MGMT* mRNA levels in HCT-116 cells and U251 cells treated with or without TMZ (1mM, 72h). Data represents means ± S.E.M. of three independent experiments. B, RT-qPCR analysis of *MGMT* mRNA levels in Ctrl and *MGMT* enhancer KO cells. Data represents means ± S.E.M. of three independent experiments.

**Supplementary Figure S3.**
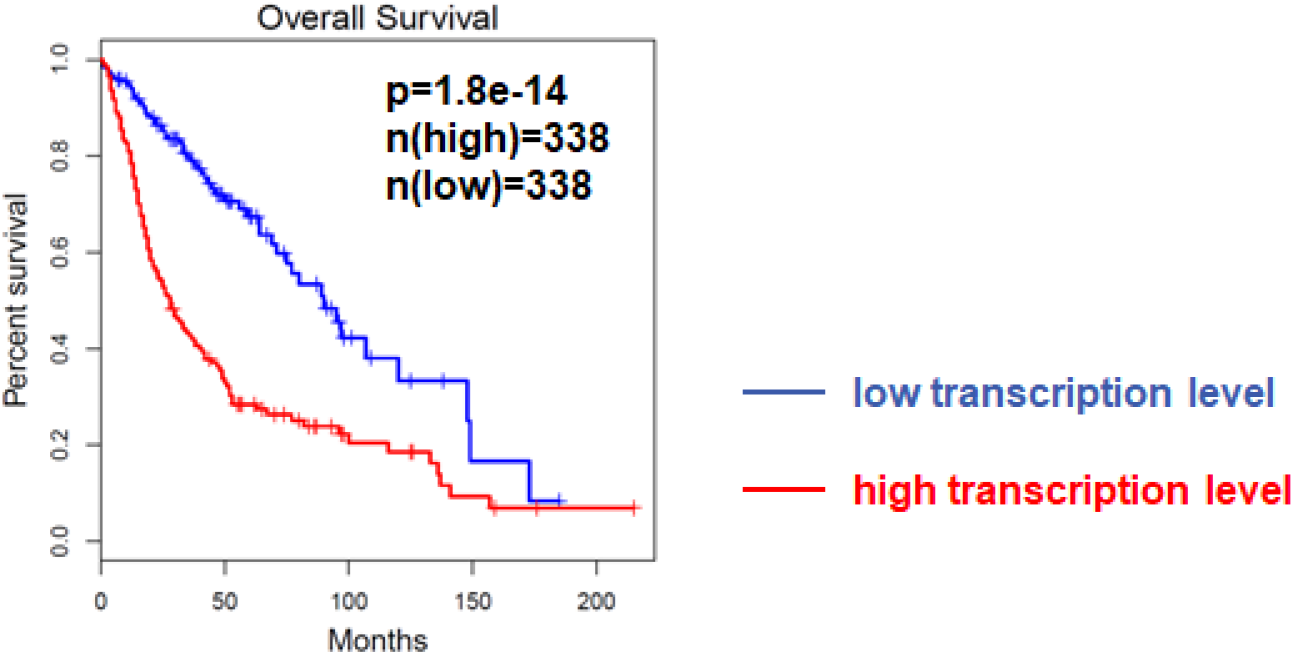
*MKI67* and associated survival data from glioma patients.

**Supplementary Table S1.**
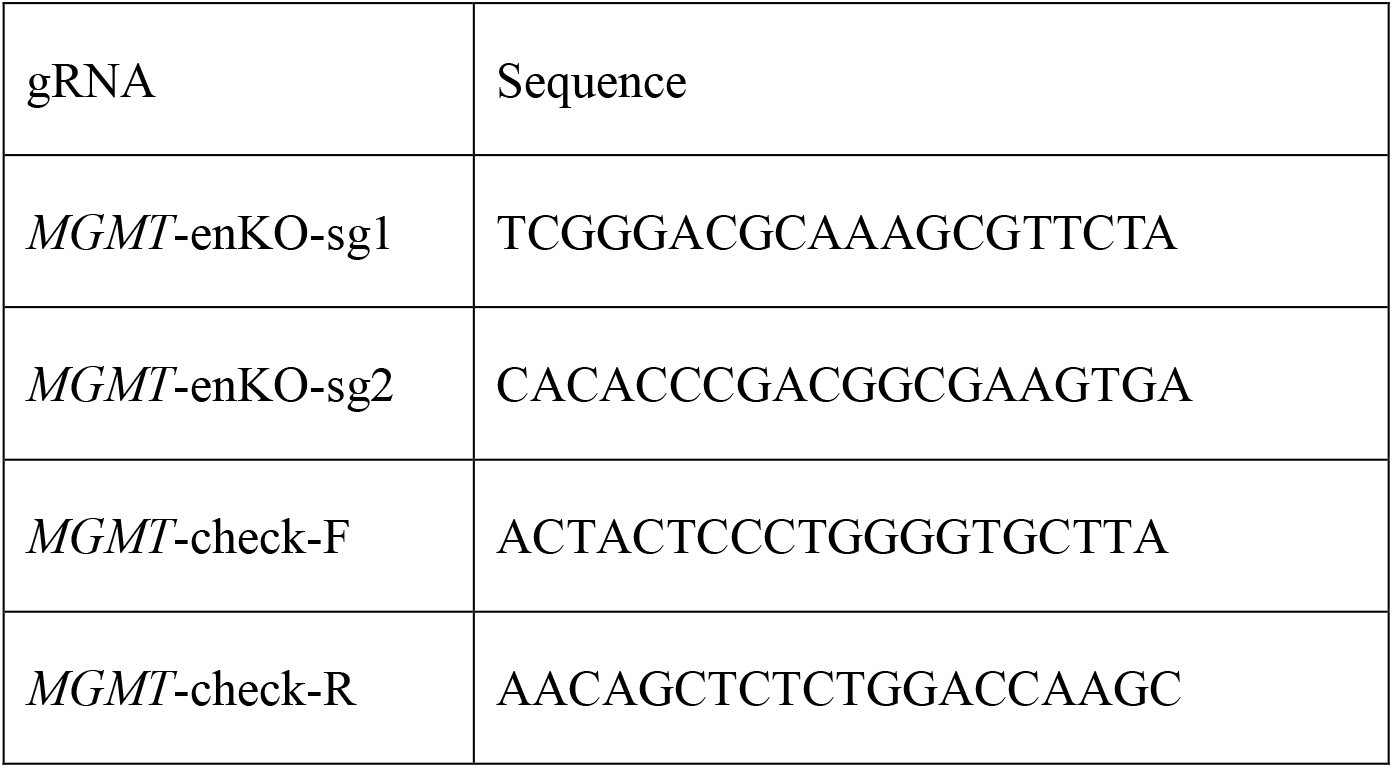
Guide RNAs and genotype primers used for CRISPR/Cas9-mediated deletions.

**Supplementary Table S2.**
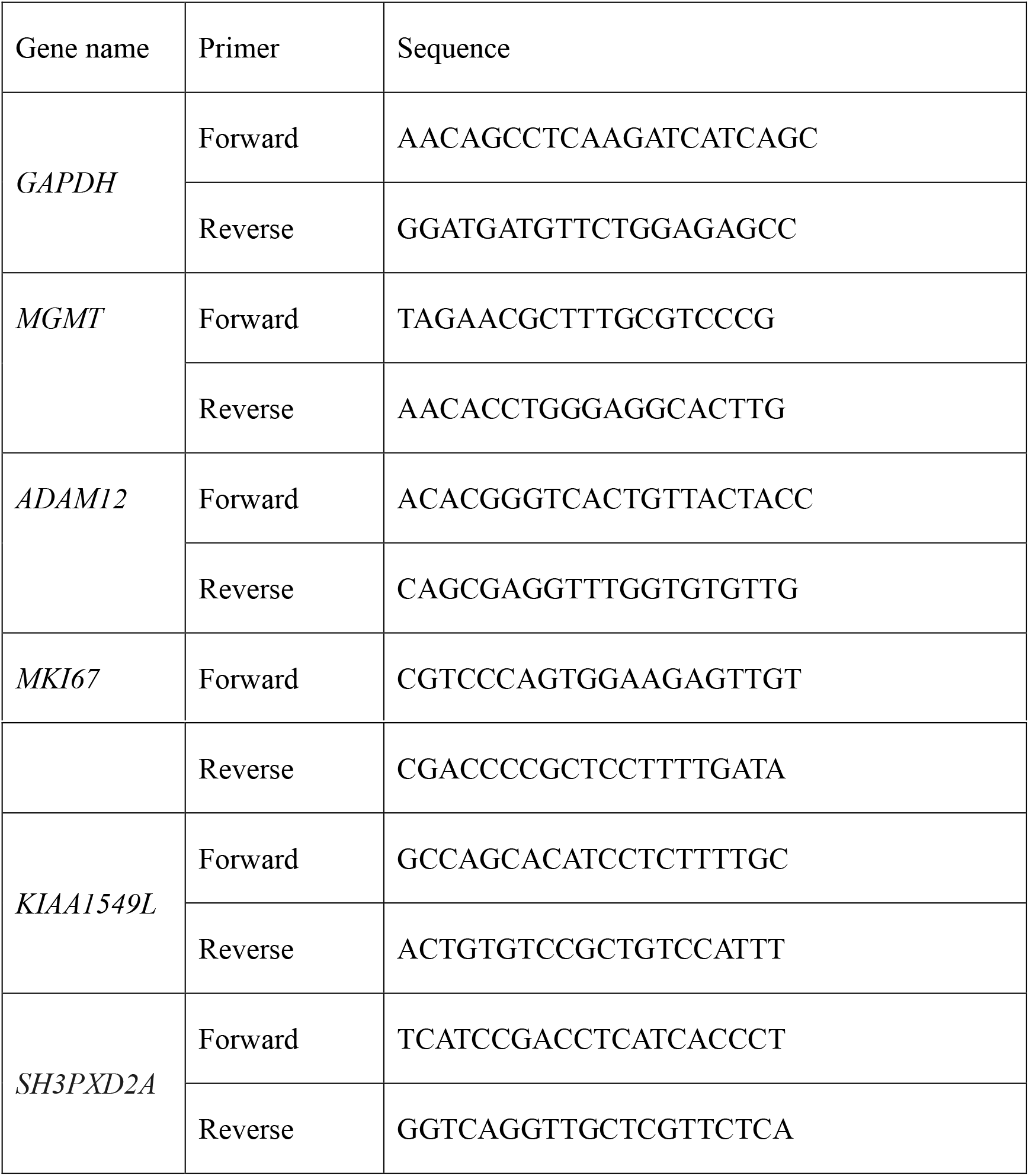
Primers used in RT-qPCR experiments.

**Supplementary Table S3.**
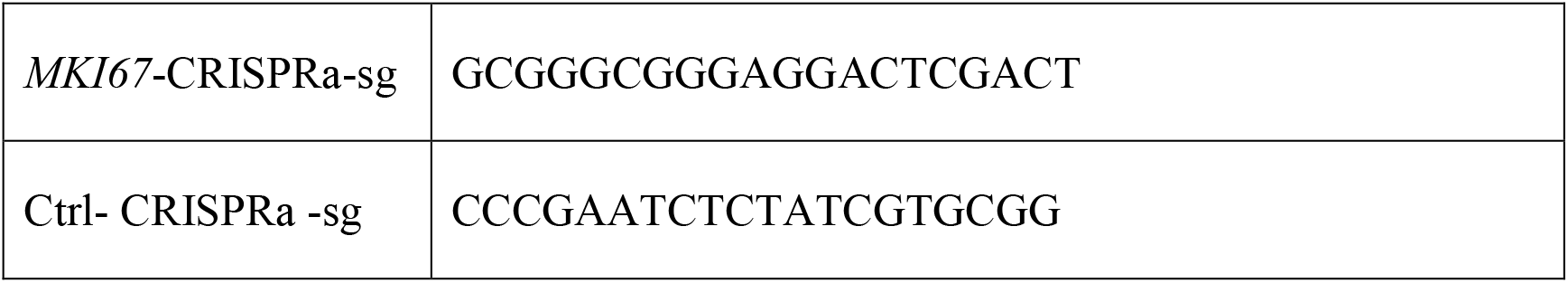
Guide RNAs used for CRISPRa.

**Supplementary Table S4.**
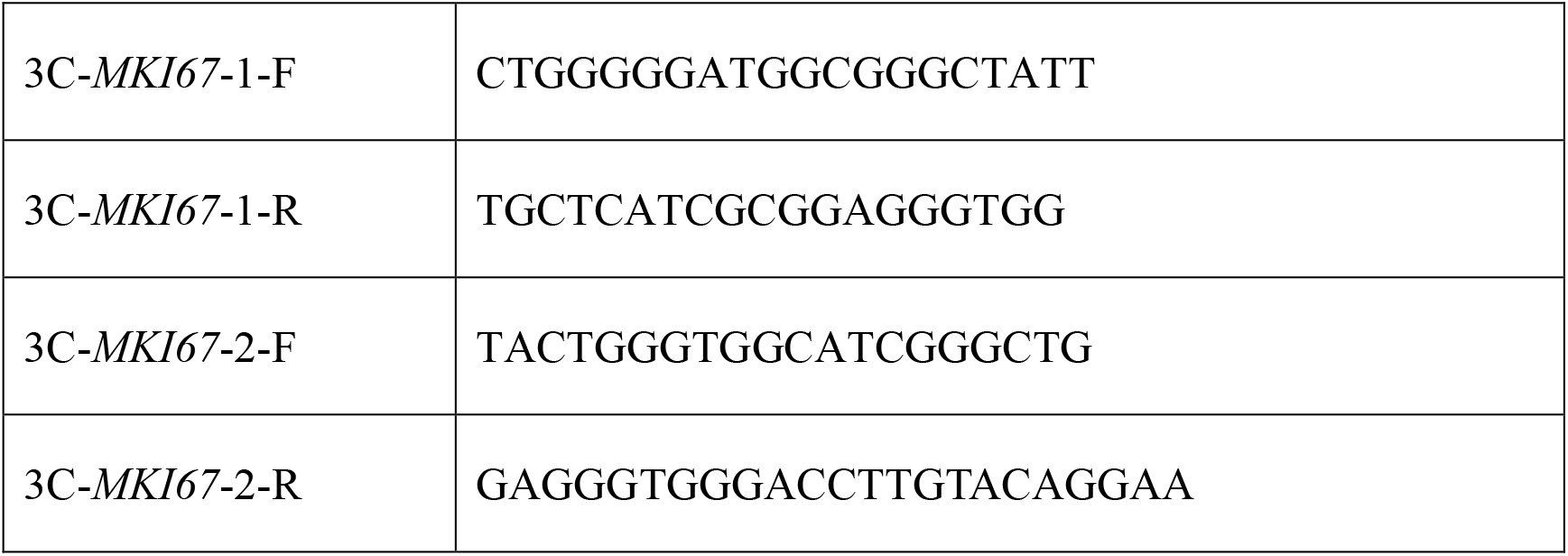
Primers for 3C-PCR assays.

